# Centrosome centering and Decentering by microtubule network rearrangement

**DOI:** 10.1101/057968

**Authors:** Gaëlle Letort, Francois Nedelec, Laurent Blanchoin, Manuel Théry

## Abstract

The centrosome is positioned at the cell center by pushing and pulling forces transmitted by microtubules (MTs). Centrosome decentering is often considered to result from asymmetric, cortical pulling forces exerted in particular by molecular motors on MTs, and controlled by external cues affecting the cell cortex locally. Here we used numerical simulations to investigate the possibility that it could equally result from the redistribution of pushing forces due to a reorientation of MTs. We first showed that MT gliding along cell edges and pivoting around the centrosome regulate MT rearrangement and thereby direct the spatial distribution of pushing forces, while the number, dynamics and stiffness of MTs determine the magnitude of these forces. By modulating these parameters, we identified different regimes, involving both pushing and pulling forces, characterized either by robust centrosome centering, robust off-centering or “reactive” positioning. In those latter conditions weak asymmetric cues can induce a misbalance of pushing and pulling forces resulting in an abrupt transition from a centered to an off-centered position. Altogether these results point at the central role played by the configuration of the MTs on the distribution of pushing forces that position the centrosome. We suggest that asymmetric external cues should not be seen as direct driver of centrosome decentering and cell polarization, but rather as inducers of an effective reorganization of the MT network, fostering centrosome motion to the cell periphery.

## Introduction

In many cells, the centrosome is positioned at the geometric center of the cell, across a wide range of conditions: in cultured cells (Burakov et al., 2003), whether they have circular or elongated shapes (Hale et al., 2011), symmetric or asymmetric adhesion patterns (Théry et al., 2006), in migrating cells (Dupin et al., 2009; Gomes et al., 2005), or in fertilized eggs (Kimura and Kimura, 2011a; Minc et al., 2011; Wühr et al., 2010). The robustness of this centering mechanism has been proposed to rely on the contribution of several types of mechanical forces acting on the MTs by pushing and pulling on cytoplasmic organelles and on cell borders, all contributing to stabilize the centrosome at the cell center (Laan et al., 2012a; Zhu et al., 2010). However, in vivo, the centrosome is mostly found at the cell periphery (Tang and Marshall, 2012). Indeed, in most differentiated cells the centrosome is anchored to the plasma membrane, where it serves as a structural base for the primary cilium (Reiter et al., 2012). This simple consideration suggests that the centrosome-MT networks not only have robust centering properties but also efficient off-centering capacities. A global understanding of MT network geometry and centrosome positioning should therefore include the striking ability of this system to easily switch from a centering to an off-centering regime.

External cues are usually considered as the main driver of centrosome decentering. Indeed, centrosome displacement to the cell periphery can be triggered by an asymmetric cue such as the contact with a neighboring cell (Rodriguez-Fraticelli et al., 2012) or a target cell (Yi et al., 2013). Such a cue generally induces local MT capture and the development of tension forces pulling the centrosome toward the cue (Combs et al., 2006; Kozlowski et al., 2007; Nguyen-Ngoc et al., 2007). However, for this mechanism to displace the centrosome up to the cell periphery, the decentering force associated to the asymmetric signal should overcome the centering forces. In such a scenario cells would have difficulties to respond to minor changes in their environment.

Here we explore the possibility that the centrosome-MT network can adopt more “reactive” conformations in which centrosome position is stable but easily destabilized by a small change in MT organization. In such a state, the centrosome is the converging point of centering and decentering forces of comparable magnitude. Therefore a mild change in an intrinsic critical parameter, or a small external cue, can be sufficient to trigger network reorganization and thus bias the force balance so that the decentering forces win over and move the centrosome to the periphery.

Minus-end directed motors, such as dynein molecules, produce pulling forces along MT length when bound to cytoplasmic vesicles or selectively on MTs tips when bound to the cell cortex (Kimura and Kimura, 2011b). The cytoplasmic localization of dyneins undoubtedly leads to a net centering force since MTs that are longer on the side of the centrosome that is facing the more distant cell edge are pulled stronger than MTs facing the closest cortex. The cytoplasmic pulling scenario also includes adherent cultivated mammalian cells with a flat (‘fried egg’) geometry, where motors, anchored on the basal surface of the cell, can pull microtubules all along their side. The contribution of dyneins anchored at the cell cortex is less clear. Cortical dyneins may have opposite effects on an isotropic radial array of MTs depending on MTs length distribution and dyneins density relative to MTs (Laan et al., 2012b). MT pushing forces, generated by MT polymerization against the cell periphery, could also center or decenter the centrosome, depending on whether MT tips can slide or not on the cortex and affect the overall network symmetry (Brito et al., 2005; Faivre-Moskalenko and Dogterom, 2002; Holy et al., 1997; Pinot et al., 2009; Tran et al., 2001). The question of centrosome positioning has been previously explored with coarse-grained models (Ma et al., 2014; Minc et al., 2011; Théry et al., 2007; Zhu et al., 2010). With this approach, individual microtubules are not represented, and the force on the centrosomes is calculated as a sum of elementary forces calculated for each angular sectors of the cell seen from the centrosome. This approach assumes that microtubules are straight, and that the ones reaching the cortex do so in the line-of-sight from the centrosome. Typically, molecular motors are also not represented, and one assumes that their contribution results either in a constant force, in the case where motors pull MT at their tip, or in a force that is proportional to the distance between the centrosome and the cortex, for motors pulling MTs on their side. Pushing forces are assumed to act purely radially, and often to be equal to the threshold for Euler-type buckling. Under these assumptions the resulting equations can be analyzed simply. In other words, MTs were generally assumed to be no longer than the cell size, and their deformation is not considered to depart from a straight configuration. This condition might hold for cells in a mitotic state but not during interphase where MTs can be much longer, and must bend to fit within the cell. This condition makes the coarse-grained approach impractical, but with modern computer hardware and state of the art simulation methods, it is possible to consider every MT explicitly, and solve the system correctly (Maly and Maly, 2010; Pinot et al., 2009; Wu et al., 2011).

To explore the balance between decentering and centering forces, and tentatively reveal some relevant cellular parameters that would be interesting to focus on experimentally, we used numerical simulations. This approach allowed us to consider the effect of several factors, which are likely to contribute to the regulation of force distribution. We examined microtubule bending and reorientation, and basic parameters, such as MT number, polymerization dynamics and stiffness, on their ability to break MT network symmetry. In this way, we identified the possible changes in network architecture that may misbalance pushing and pulling forces and promote centrosome decentering.

## Results

Centrosome positioning mechanisms are challenging to study since numerous factors such as cell shape, MTs properties or interacting proteins intervene and vary in different cell types and experimental conditions. Furthermore, there is currently no experimental way to measure the mechanical forces experienced by MTs *in vivo*, precluding the mapping of the spatial distribution of pushing and pulling forces that can be used for centering. Here, simulations were performed with the software Cytosim (Nedelec and Foethke, 2007). This cytoskeleton simulator is based on Langevin dynamics approach and offers the possibility to take into account numerous components in minimal computational time thanks to a semi-implicit numerical integration scheme (Kozlowski et al., 2007; Loughlin et al., 2010, 2011; Ward et al., 2014).

We simulated pure centrosomal arrays, in which all MTs are attached to a common center at their minus ends. The angular distribution of MT nucleation was isotropic. We simulated systems containing one centrosome composed of 100 MTs for 400 s. MT growth followed the classical two-states dynamic instability model (see supplementary table 1 and Material and Methods for all numerical parameters). MTs were confined to regular geometries representing different idealized cell shapes. They could bend as linear elastic beams, and thus follow Euler's buckling theory. Entities that could bind/unbind and move along MTs were added to simulate the action of minus end directed motors. Centrosome displacement is opposed by a viscous drag calculated to match the experimental observations. MTs growing against geometrical boundaries produced pushing forces whereas minus-end directed motors generated the pulling forces. By simply monitoring the position of the centrosomes, we could deduce if the tested conditions resulted in a net centering or a decentering effect.

## Contribution of pulling forces

MTs generate pushing forces as they polymerize against a barrier (Dogterom and Yurke, 1997). The spatial distribution of growing MTs within the cell determines the net force transmitted to the centrosome. If the aster is isotropic, pushing forces are directed toward the center of the cell, but they can be directed away from it in the case of anisotropic distribution (Pavin et al., 2012). Therefore, any parameters influencing MTs spatial distribution, such as nucleation, dynamics or forces that induce bending are likely to strongly affect the impact of pushing force on the centrosome. MTs are in particular easily deflected by forces applied on their ends. The net force on the centrosome will depend on MT stiffness, the number of MTs and their configurations.

To further investigate the effect of the cellular geometry on centrosome positioning, we switched to ellipsoidal, rectangular and triangular geometries. We explored the two fundamental motor distributions, systematically varying the total number of motors in the system to explore different ratio of motors to MTs. In the case of cytoplasmic localization of dyneins, the centrosome moved toward the center of gravity of the shape for any given initial position and all tested geometries (Figure 2A), consistent with experimental observation in sea urchin eggs (Minc et al., 2011). In contrast, in the case of a cortical distribution, centrosomes did not move toward the center of gravity in the triangular geometries. After fast and erratic displacements throughout the cell, the centrosome usually converged toward the middle of the longest edge, which contains more dyneins (Figure 2B, C). Like in the circular geometry, switching to a cortical distribution led to stronger net forces as illustrated by faster centrosome displacements (Figure 2C). These results showed how uneven angular distribution of cortical dyneins can act as a strong decentering force whereas in the case of cytoplasmic dyneins the net force was weaker and systematically directed toward the cell center.

**Figure 1.**
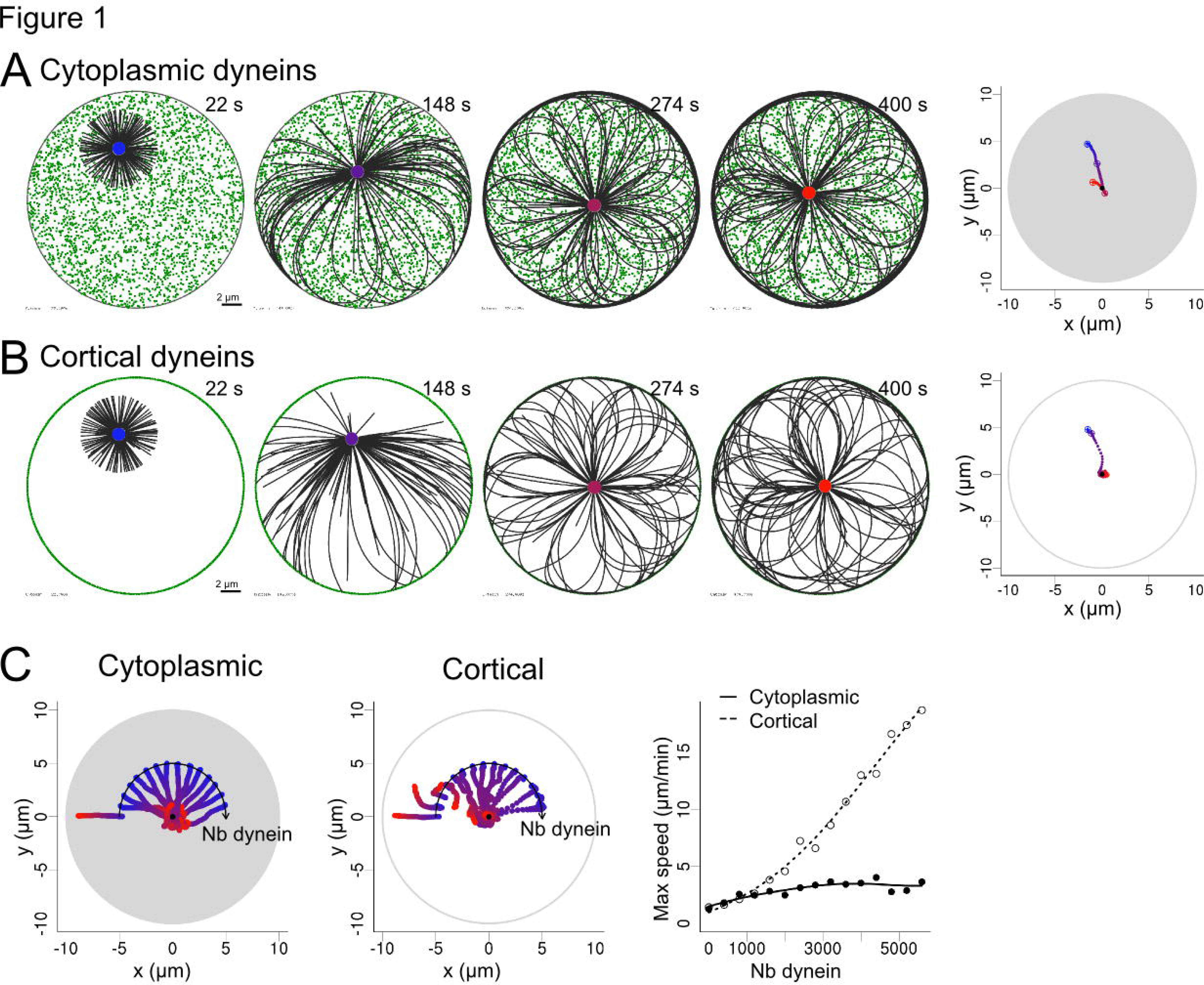
Centrosome centering by pulling forces. (A) Simulation where the motors are distributed in the cell. (B) Simulation where the motors are distributed only on the edge the cell. Dynein motors are shown in green and MTs in black. The centrosome positions are indicated by colored points, from blue (0 s) to red (400 s). (right most column) Trajectory of the centrosome in the simulation shown on the left panel. The grey shade represents the area that is filled by motors. (C) Variation of the number of motors for both cytoplasmic and cortical motor distribution. 15 trajectories are shown on each plot, where the number of simulated dyneins is increased from 0 to 7000 with a step of 500. The initial position of the centrosome was set on an arc axis to make them all visible on a single plot. This should not affect the outcome of the simulation, since the system has rotational symmetry. In all case, the centrosomes are initially placed at a distance from the center corresponding to half the cell radius. (Right) Maximal speed reached by centrosome during each simulation as a function of the number of dyneins, for both cortical and cytoplasmic distributions. Each symbol is the results of one simulation, and the line is a guide for the eye.

**Figure 2.**
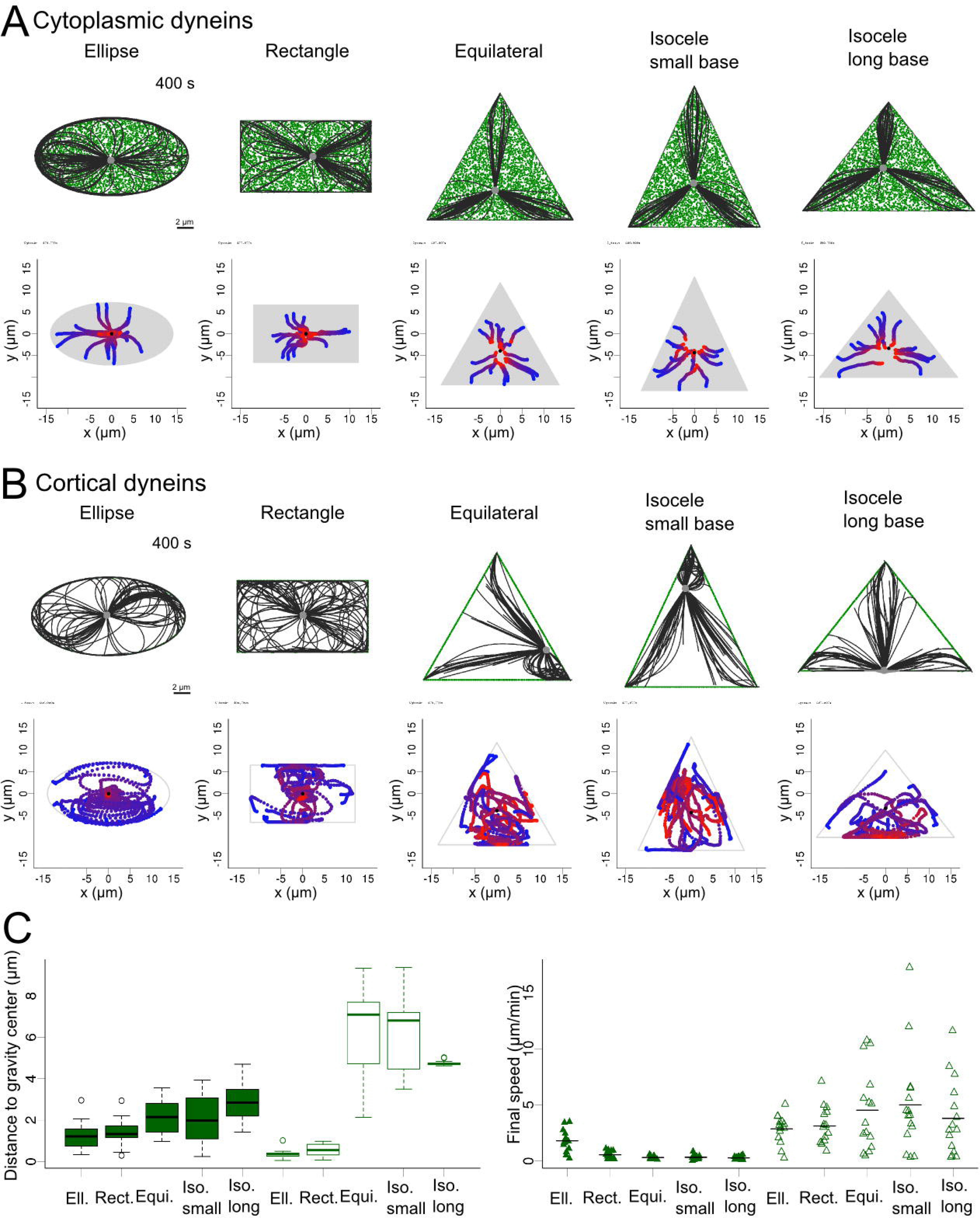
Centrosome positioning by pulling forces. (A-B) (top) Snapshot after 400 s of a centrosome simulated in different geometries: ellipse, rectangle, equilateral triangle, acute isosceles triangles and isosceles triangle whose base is the longest side. Dynein are shown in green, MTs in black and centrosome in gray. Motors are either distributed (A) cytoplasmically or (B) cortically. (Bottom) Trajectories of centrosomes in 15 simulations for each geometry. Centrosome position is indicated by colored points, from blue (0 s) to red (400 s) in the trajectories plots. Gray area represents the area covered by motors. Black dot indicates the center of gravity of the shape. (C) (left) Boxplot of the distance of centrosome to gravity center after 400s for each geometries, for cytoplasmic and cortical distributions (full and empty boxes, respectively). (right) Stripchart of the final speed of centrosome in the simulations for all geometries for cytoplasmic and cortical distributions (full and empty triangles, respectively).

## Contribution of pushing forces

MTs generate pushing forces as they polymerize against a barrier (Dogterom and Yurke, 1997). The spatial distribution of growing MTs within the cell impact the direction of the net pushing force exerted on the centrosome. If the aster is isotropic, pushing forces are directed toward the centrosomes, but they are directed away from it in the case of anisotropic distribution (Pavin et al., 2012). Therefore, any parameters influencing MTs spatial distribution, such as nucleation, dynamics or forces that induce bending are likely to play key roles in determining the direction of the net pushing force on the centrosome. MTs are in particular easily deflected by forces applied on their ends. The net force on the centrosome will depend on MT stiffness, the number of MTs and their configurations.

To investigate these effects, we considered a radial array of flexible MTs with their minus-end anchored at the centrosome. MTs are anchored at regular angular intervals, such as to form an isotropic aster, but we tuned the angular stiffness of their anchorage, to allow them to pivot at various degrees around their minus-end anchorage. In the basic setup, plus-ends could glide along the edge of the cell, as the contact is considered to be frictionless, but by adding immobile anchors at the cortex, that capture the plus-ends and pin them we can also prevent sliding. Thus varying the angular stiffness at the centrosomes, and the number of anchors points at the cortex, enabled us to test the combinatorial effects of allowing or disabling minus-end pivoting and plus-end gliding (Figure 3A). At first, those simulations were performed in the absence of minus-end motor associated pulling forces. As expected from MT observation in lymphoblastic cell lines (Bornens et al., 1989) and previous numerical simulations (Pinot et al., 2009), when both pivoting and gliding were allowed the network became asymmetric and pushed the centrosome off-center toward the closest edge (Figure 3A, top left). Indeed, MTs oriented toward the closest side were the first to reach the cortex and glide toward the opposite direction to minimize the bending energy associated to their curvature. This effect was reduced if MT pivoting was forbidden, as the aster remained more isotropic (Figure 3A, top, right). Strikingly, when MT gliding along the cell cortex was prevented, those first MTs in contact with the cell cortex were pinned and pushed on the centrosome, which got displaced toward the opposite cell edge (Figure 3A, bottom left). MT pivoting ability allowed them to join and form comet-like tail pushing the centrosome as observed in *Dictyostelium* (Brito et al., 2005). Interestingly, when both gliding and pivoting were prevented the network never became asymmetric. Instead, it rotated briefly and adopted a vortex-shaped conformation in which the pushing forces kept the centrosome near the cell center (Figure 3A, bottom left). This centering effect by symmetric pushing forces was quite robust and independent of cell geometry (Figure S1). As we reduced the strength of either the pivoting or the gliding stiffness while fixing the other, the network displayed rapid transition from centering to decentering (Figure 3C). This suggested that the modulation of these parameters can be an efficient way to break network symmetry and place the centrosome near the cell periphery (Figure 3B and 3C). Importantly, these behaviors were independent on centrosome initial position (Figure 3B), meaning that if the centrosome is initially positioned at the cell center, the symmetry in the network will be spontaneously broken either because MT pivot, glide or both, leading to centrosome decentering.

**Figure 3.**
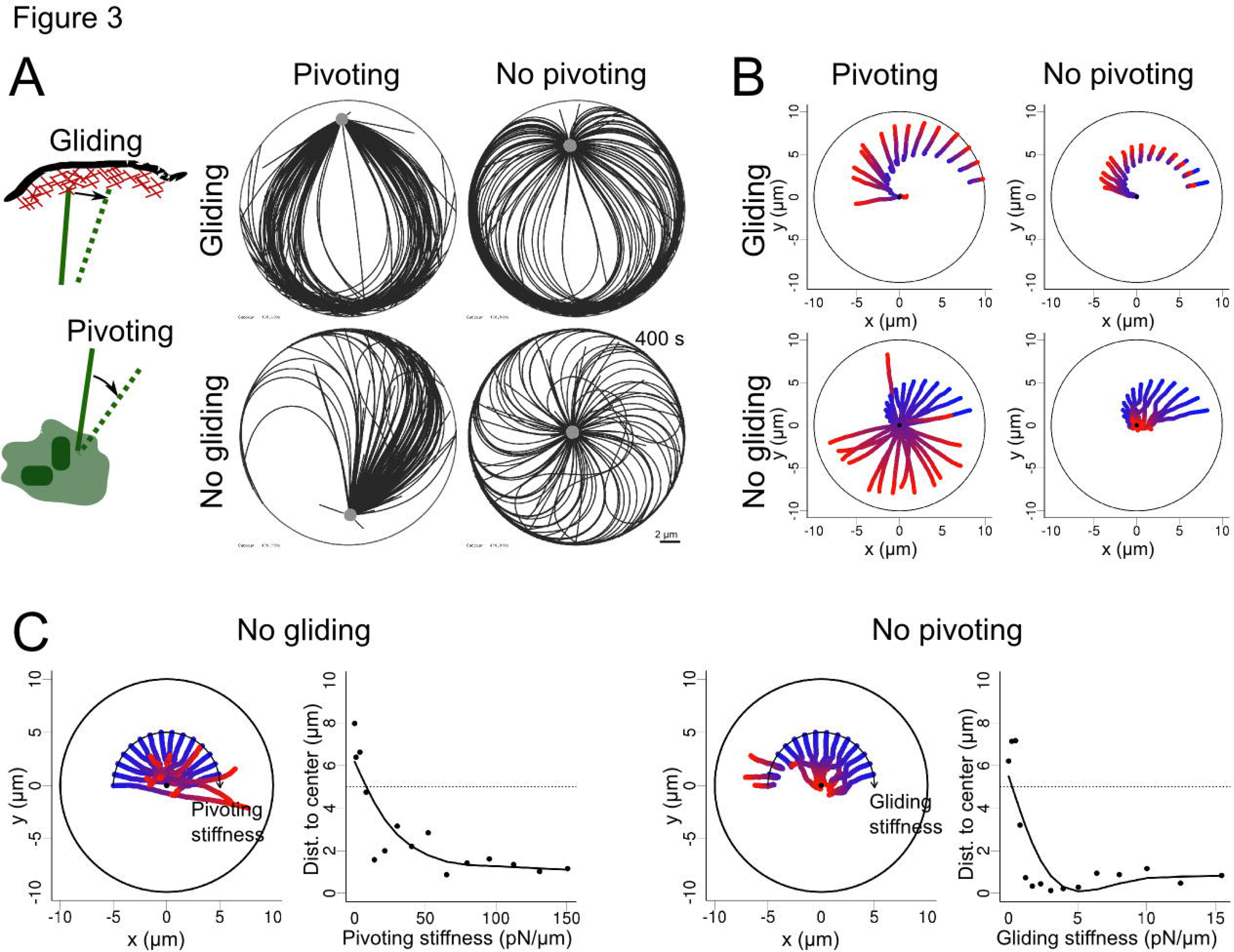
MT network rearrangement in the presence of pushing forces. (A) (left) schematic representation of a MT plus end gliding at the cortex, and of a MT pivoting around its anchor point in the centrosome. MTs and centrosomal complex are in green, actin cortex in red. (right) Snapshots of simulations (400 s) covering all the possibilities, obtained when pivoting and gliding are independently allowed or not. (B) Trajectories of centrosomes in 15 simulations where the centrosome was initially positioned at different distance from the cell center, also for each of the pivoting/gliding conditions. The center of each disc is marked with a black point. (C) (left) Trajectories of centrosomes in simulations with varied pivoting stiffness when gliding is not allowed. Pivoting stiffness is varied from 0 to 150 pN/pm from left to right along the arc. (right) Centrosome positioning as a function of pivoting stiffness, measured by the distance to the cell center. The dashed-line indicates the initial centrosome-center distance. (D) (left) Trajectories of centrosome in simulations with varying gliding stiffness when pivoting is not allowed. Gliding stiffness is varied from 0 to 15.5 pN/pm from left to right along the arc. (Right) Centrosome positioning as a function of gliding stiffness, measured by the distance to the cell center. Dashed-line indicates the initial centrosome-center distance. (A-D) The colors of the centrosome trajectories indicate time, from blue (0 s) to red (400 s) in all the plots.

We then studied how MT stiffness, number and length affected these behaviors. In the symmetric conformations of the network, when both MT gliding and pivoting were restricted, centering appeared robust to a decrease in MT number and length (Figure S2). The effects of changing MT stiffness were mild, between 20 and 100 pN/μm2. Very soft MTs could not transmit polymerization forces efficiently, whereas very rigid MTs could not bend after reaching the cortex, in both cases freezing the centrosome motion (Figure S2). If the MT network lost its isotropy, either because MT gliding along edges or pivoting around the base are allowed, then increasing the number of MTs progressively reinforced the net pushing forces and decentered the centrosome (Figure 4). We observed this effect either by raising the number of nucleation sites or by decreasing the catastrophe rate. Increasing the stiffness of MTs also produced the same outcome. All these parameters are therefore interesting targets if one wants to induce centrosome decentering.

**Figure 4.**
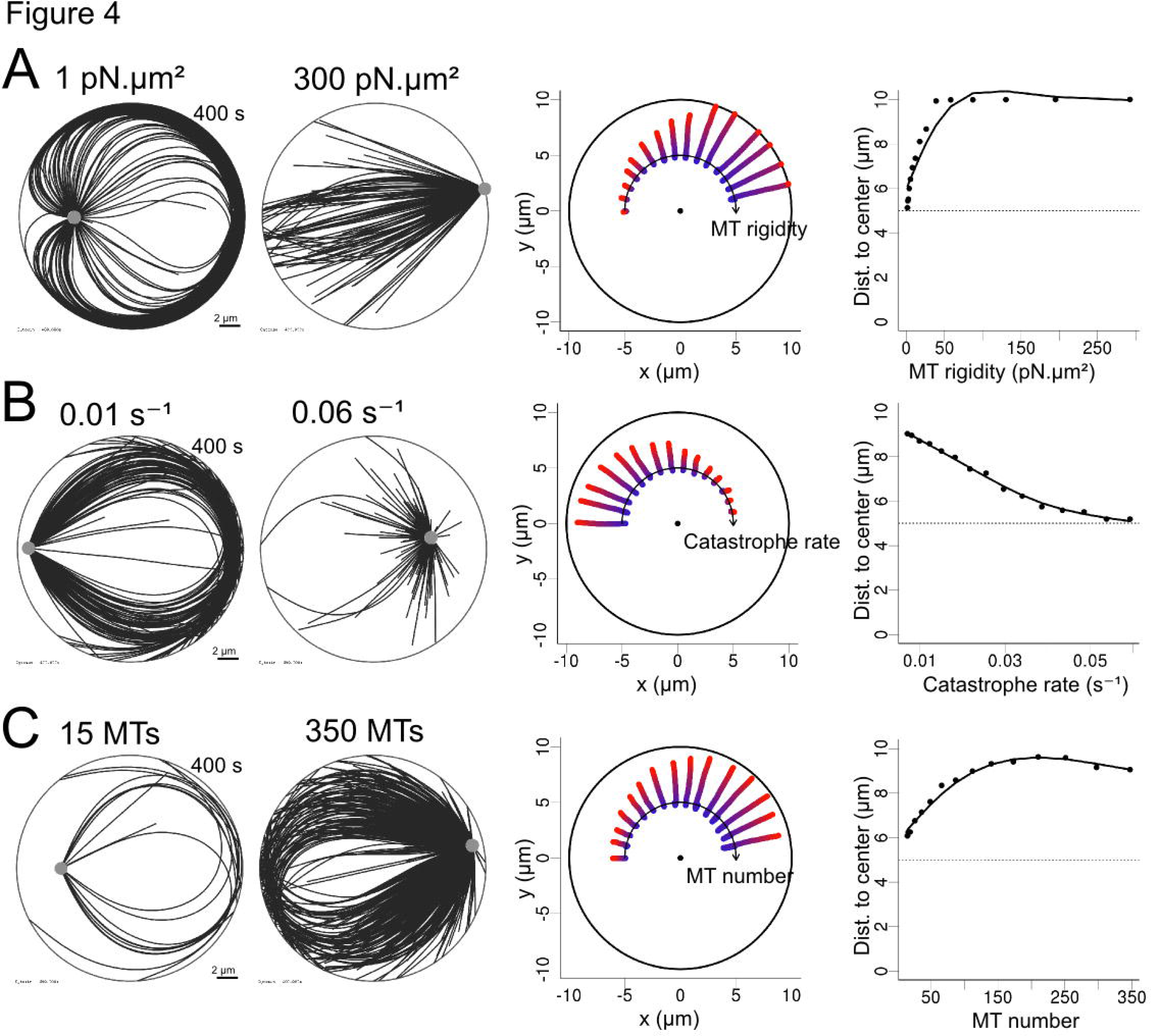
Efficiency of pushing forces (with pivoting and gliding allowed). (A-C) Simulations where one parameter was varied: MT rigidity, from 1 to 300 pN.μm^2^; of MT unloaded catastrophe rate from 0.01 to 0.06 /s; and of the number of MT in the aster, from 15 to 350. (left column) Two exemplary simulations (400 s) obtained by varying one parameter in each case. (middle column) Trajectories of centrosomes obtained by varying one parameter, displayed along an arc, with increasing values from left to right. For each trajectory, time is indicated by the color, from blue (0 s) to red (400 s). The center of each disc is marked with a black dot. (right) Centrosome positioning as a function of various parameter, measured by the distance to the cell center. The dashed lines represent the initial centrosome-center distance (half the confinement radius).

### Transitions from centering to decentering regimes

We then combined the pulling forces due to minus-end directed motors and the pushing forces due to MT polymerization to investigate the potential transition from centering to decentering regimes. Since dynein inhibition has been shown to induce centrosome decentering (Burakov et al., 2003; Wu et al., 2011), we assumed that dynein-associated forces contributed to center the centrosome, i.e. that dynein molecules are anchored in the cytoplasm rather than at the cell cortex. We first wanted to know if a centrosome could spontaneously move off center in the absence of asymmetric cues. We thus tested if a global variation of the MT network properties, such as the parameters described above, could overcome the centering effect of cytoplasmic dyneins and promote centrosome displacement to a peripheral position.

As described in the first part of this study, with high concentration of cytoplasmic dyneins (4000 dyneins per cell corresponding to 40 per MT), MT outward pushing forces were not sufficient to overcome the dynein-induced centering effect (Figure 1, 2). Transitions were seen only in cells where the dynein concentration was reduced. In the following simulations we used 100 dyneins per cell (ie 1 per MT). This condition is physiologically relevant since it has been shown that pushing and pulling forces are of comparable magnitude (Zhu et al., 2010). We thus studied the conditions in which variations in MT rigidity, number and catastrophe rate, as well as centrosome pivoting and cortex gliding stiffness, could lead to decentering despite the dynein-induced centering pulling forces.

As MT rigidity was increased, the net force applied on the centrosome progressively switched from centering to decentering (Figure 5A). However, variations of catastrophe rate and number of MTs, although able to increase pushing forces magnitude (Figure 4), did not cause decentering (not shown). This is due to the magnitude of the forces applied to MTs tips, which are such that they induce catastrophe events and thus limit the efficiency of pushing force. However, transitions from centering to decentering, was observed in smaller cells (of radius 7 μm) where the pushing forces are stronger (because they scale in 1/L^2^ according to Euler's theory) (Figure 5B, C). Reducing drastically the catastrophe rate turned the centering regime into a weak decentering one (Figure 5B). The system was quite sensitive to the number of MTs, and a progressive transition from centering to decentering occurs as the number of MTs increased (Figure 5C). Varying either centrosome angular stiffness (which affected pivoting) or cortex anchor stiffness (which affected sliding) had even more drastic effects, and could induce abrupt transitions in the position of the centrosome (Figure 5D, E). From these results, the ability of the network to reorganize asymmetrically appears as a key regulator of the force balance at the centrosome.

**Figure 5.**
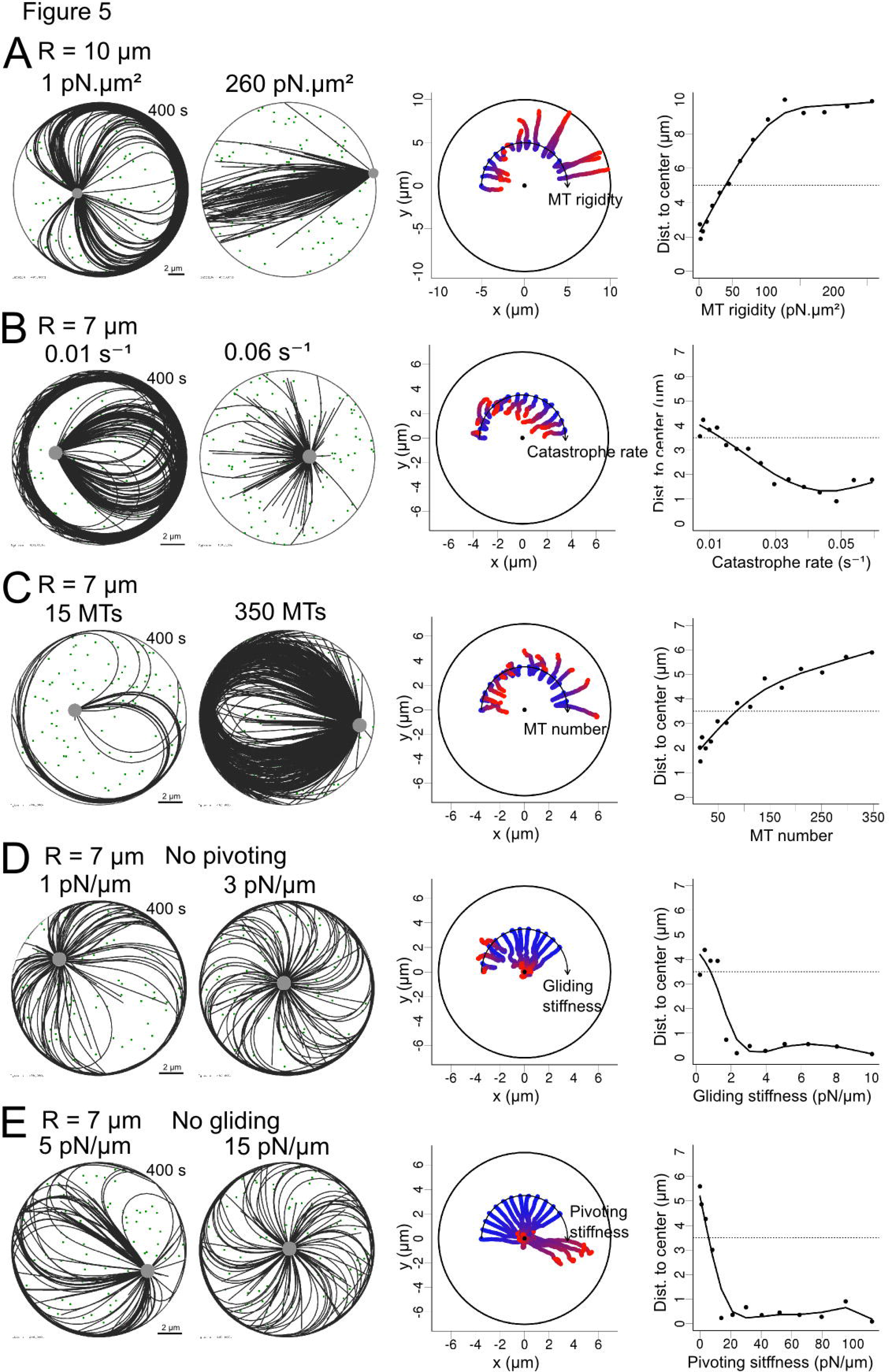
Transitions between centering and decentering regimes. (A-E) Simulations where one parameter was varied: MT rigidity, from 1 to 260 pN μm^2^; MT unloaded catastrophe rate from 0.01 to 0.06 /s; number of MTs from 15 to 350; MT gliding stiffness while pivoting is not allowed, from 0 to 10 pN/pm and MT pivoting stiffness when gliding is not allowed, from 0 to 110 pN/pm. In each case, 100 cytoplasmic dynein motors are randomly positioned in the cell, and one parameter of the system is varied systematically. Cell radius is 10 pm in (A) and 7 pm in (B-E). (Leftmost two columns) Exemplary simulations for 2 different outcomes observed while varying a parameter. (middle column) Trajectories of simulations obtained while varying a parameter, and displayed along an arc, with values increasing from left to right. For each trajectory, time is indicated by the color, from blue (0 s) to red (400 s). The center of each disc is marked with a black dot. (right column) Centrosome positioning as a function of various parameter, measured by the distance to the cell center. Dashed-line represents the initial centrosome-center distance (half the confinement radius). (A-C) MTs are allowed to pivot and glide.

Noteworthy, these results showed that moderate changes in any of the tested parameters, isolated or in parallel, were often sufficient to induce a complete reversal of situation where the centrosomes is located either at the center or near the periphery of the cell. In the case where the transition is not complete, the parameter change could anyway contribute to prime the network for such a transition. For example, allowing the network to be asymmetric increased the net force on the centrosome. Even when this force is effectively balanced by the inward pulling forces exerted by cytoplasmic dynein, it made the reversal more likely to occur if another perturbation is added. For example, the net inward pulling force can be further reduced by additional asymmetric outward pulling forces, such as those exerted along cell-cell contacts. This idea was tested by the addition of external cues, which were simulated by adding cortical dyneins within a 60° wide crescent. We then compared the response of a “constrained” network, in which MT central pivoting and peripheral gliding were forbidden, and the response of a “reconfigurable” network in which pivoting and gliding are allowed, to increasing amounts of localized cortical dyneins (Figure 6A). The constrained network was less sensitive to asymmetric pulling forces since the pushing from the captured MTs opposed the tension developed by the cortical dyneins (Figure 6A). By contrast, the reorientation of MTs in the “reconfigurable” network, redirected some polymerization forces toward the cortical pulling site (Figure 6B). Thus, the “reconfigurable” network appeared more responsive to external cues, and the centrosome is decentered more readily (Figure 6C). Together these results indicate that several intrinsic parameters of the MT networks such as MT number, rigidity and in particular the ability to reorient MTs is key to modulate the response of the cell. The magnitude and distribution of pushing forces could either lead to robust centering of the centrosome or place it in a reactive conformation, where it can respond to weaker external asymmetric stimulation. Such reactive conformations are characterized by spontaneous symmetry breaking events and centrosome decentering in the absence of asymmetric external cues.

**Figure 6.**
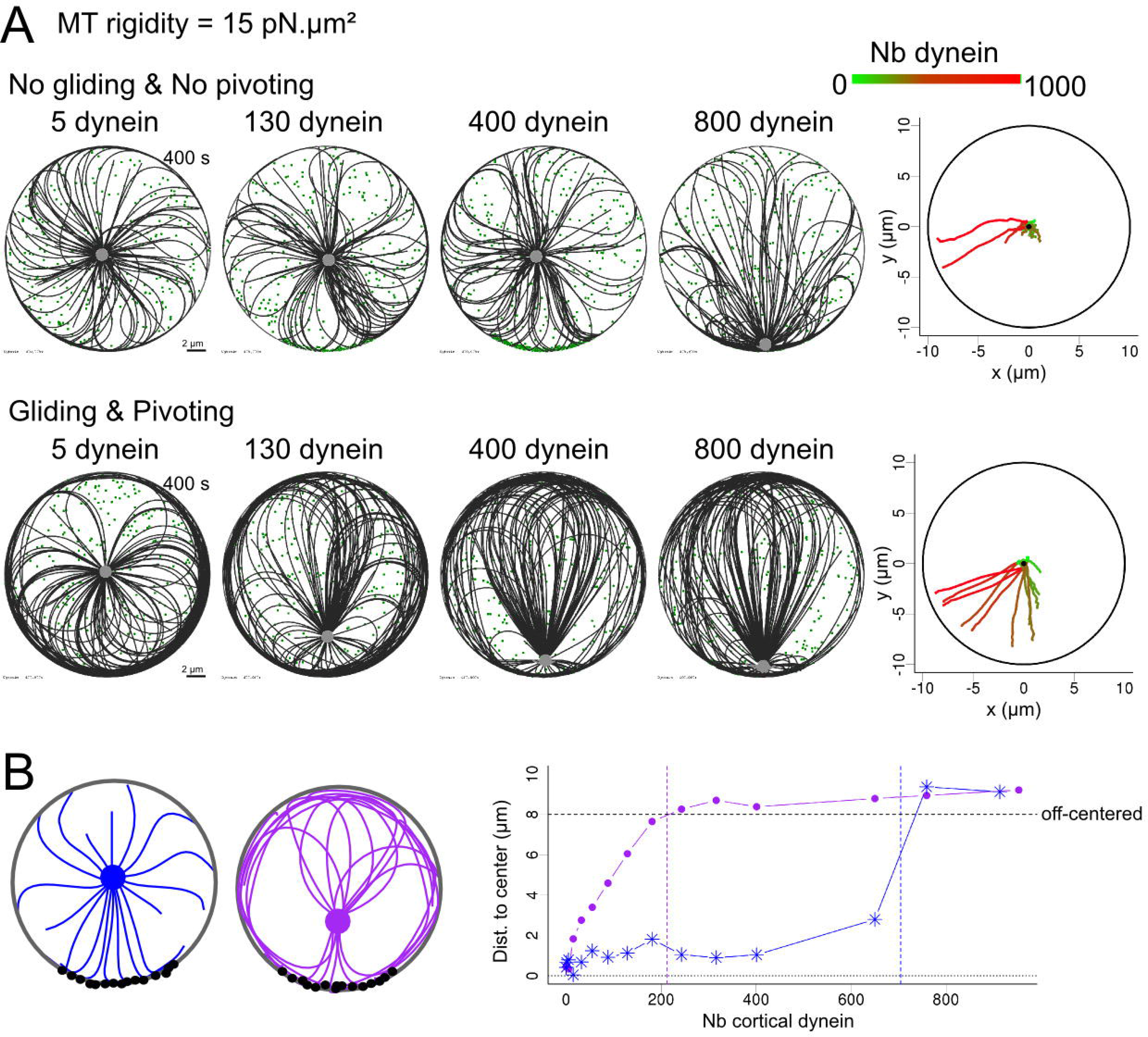
Sensitivity of centrosome positioning to external cues modulated by internal properties. (A-B) The centrosome is initially placed in the center of the cell and is decentered. The cell has a radius of 10 pm and contains 300 randomly positioned cytoplasmic dyneins. MT rigidity is set to 15 pN μm^2^. (A) Simulations where MT pivoting and gliding are not allowed (top) and where MT pivoting and gliding are allowed (bottom). (Left) Simulations (400 s) in which different number (5, 130, 400 and 800) of cortical dynein have been added on a 60° crescent at the “bottom” part of the cell. (Right) Trajectories of centrosomes, in a color representing the number of dynein in the cell from 0 (green) to 1000 (red). The position of the cortical crescent was shifted to be able to distinguish the different trajectories on a single plot. (B) (Left) schematic representation of MT network configuration when MT pivoting and gliding are not allowed (blue) and when both are allowed (purple). The cortical dynein molecules are represented in black. (Right) Final distance of centrosome to cell center according to the number of cortical dynein placed on the crescent, when gliding and pivoting are not allowed (blue) and allowed (purple). The black horizontal dashed line indicate the threshold above which we considered the centrosome to be off center. The colored vertical dashed lines represent the threshold number of motors necessary to be decentered, in each case.

## Discussion

The process of centrosome positioning is still under investigation, in particular because different mechanisms can prevail in different cellular conditions (Ma et al., 2014). Here we focused on conditions in which MTs can be longer than the cell diameter, which is the case for adult mammalian cells in interphase (Wu et al., 2011) rather than the more usually studied large mitotic cells in embryos (Kimura and Onami, 2007; Minc et al., 2011; Wühr et al., 2010). In this condition, several parameters could regulate the symmetry of MT network architecture, independently of external cues or preexisting asymmetries in boundary conditions such as local capture, stabilization or mechanical forces. MTs number, length and rigidity as well as centrosome stiffness all have the ability to induce a spontaneous network symmetry break and thus lead to centrosome decentering.

Centering capacities are generally considered to result from minus-end-directed motors such as dyneins (Kimura and Kimura, 2011b). Here, by using numerical simulations, we compared the contributions of cytoplasmic and cortical dyneins. We confirmed that cytoplasmic dynein molecules induce robust centrosome centering, whereas cortically anchored dyneins are capable of promoting either centering or decentering depending on cell shape (Ma et al., 2014). Asymmetric pulling forces due to cell shape extension promoted centrosome decentering toward the cell longest edge.

Next, we studied how the MT network configuration determined the orientations of pushing forces associated with MT polymerization. When MTs were tightly constrained in space, i.e. if they cannot pivot around their anchor point in the centrosome and if their plus-ends may not glide along the cell edge, they maintain an isotropic distribution and generate centering forces. In contrast, when the MTs are free to pivot or slide, the net pushing force pushes the centrosome toward the periphery. In both scenarios, MTs' stiffness and number could modulate the speed at which the centrosome moves. Thus when pushing forces were opposed by dynein induced pulling forces, modifying the number of MTs or their stiffness could trigger a transition from a regime in which the centrosome is at the center, to a regime in which the centrosome adopts a peripheral position.

We investigated some of the theoretical mechanisms that can affect MT architecture and centrosome positioning, limiting our approach to numerical experiments. The results pointed at the possible role of several parameters, suggesting experimental investigations.

Centrosome angular anchor stiffness appeared as a critical parameter since small variations from 20 to 5 pN/μm were sufficient to induce an abrupt transition from a centering to a decentering regime (Figure 5E). The anchoring of MTs at the centrosome is not well characterized. MTs can detach from the centrosome in mammalian cells (Alieva et al., 2015; Keating and Borisy, 1999) and were seen to pivot around yeast mitotic spindle poles (Kalinina et al., 2012), but the angular stiffness has not been measured. The pivoting of a MT with respect to the centrosome is likely to depend primarily on the stiffness of the pericentriolar material in which MTs are embedded. Regulation of pericentriolar material density and crosslinking density (Woodruff et al., 2014), as well as the polymerization of actin filaments nearby (Farina et al., 2015), could possibly affect this stiffness. Moreover, the ability of MTs under tension to tear apart pieces of pericentriolar material during specific phases of the cell cycle suggests that the material stiffness is precisely regulated (Krueger et al., 2010; Megraw et al., 2002; Rusan and Wadsworth, 2005).

Pushing forces naturally scale proportionally with the number of MTs in the aster, and scale inversely with the length of MTs, but these parameters also have a less obvious effect on the network symmetry (Figure 4). These parameters vary widely from one cell type to another, and also vary during cell cycle progression (Piehl et al., 2004). It would be interesting to look at them in more details during centering to decentering transitions, for instance during epithelial morphogenesis, ciliogenesis or immune synapse formation.

Our simulations confirmed a strong role for MT stiffness, which was expected since MT bending stiffness is a key parameter to the transmission of MT polymerization force produced at the plus-ends to the centrosome attached at the minus-ends and thereby regulates the net force on the centrosome. Increasing the MT stiffness is sufficient to switch from a centering to a decentering regime (Figure 5A). Several parameters have been shown to affect MT rigidity (Hawkins et al., 2010). MT associated proteins that can either increase (Felgner et al., 1997) or decrease (Portran et al., 2013) MT bending stiffness. MT crosslinking molecules can also modulate the size of MT bundles, and consequently their rigidity (Bathe et al., 2008) and thus affect the system similarly.

Cortical stiffness in our simulations reflects the interaction that MTs have with the cell cortex, and is a property of the cell cortex. MT ability to glide or not along cell cortex was key to network rearrangement, to symmetry breaking, and to the orientation of pushing forces toward the cell periphery (Figure 3, Figure 5D). This parameter reflects that MTs could get entangled into a crowded cortical actin network or be physically linked to those microfilaments (Coles and Bradke, 2015). Plus-ends have been seen to grow or not along cell periphery depending on the presence of filament bundles or branched meshwork suggesting that the local actin architecture could regulate MT gliding (Théry et al., 2006). Accordingly, changes in cell adhesion and modifications of the associated cortical actin network could result in MT network rearrangements and finally induce centrosome repositioning.

For the sake of simplicity, several factors were not taken into account in our simulations, notably the contribution of non-centrosomal MTs (Alieva et al., 2015) and kinesins (Cross and McAinsh, 2014) although we know that these factors contribute to the intrinsic regulation of force production and network reorganization. We also ignored key external contribution such as centrosome and MT interactions with the nucleus (Burakov and Nadezhdina, 2013) and the production of forces by the actin cytoskeleton (Gupton et al., 2002; Waterman-Storer and Salmon, 1997). Importantly as well, we have considered that the cytoplasm was devoid of obstacles, and that MT motions were only hindered by the cell boundaries. The elasticity of the actin network and other obstacles surrounding MTs may however constrain their deformation and significantly increase the magnitude of pushing forces transmitted to the centrosome (Brangwynne et al., 2006; Shan et al., 2012). These important parameters deserve further investigation.

In this work, we studied the ability of MT asters to become anisotropic, a reorganization of the MT network that ultimately pushed the centrosome near the cell periphery. In physiological conditions the MT cytoskeleton within a cell is rarely isolated as cells contact other cells. These contact represent external cues that affect the MT network within each cell. A MT network spanning the entire cytoplasm can physically integrate all these contributions from the surrounding tissue. Ultimately, the position of the centrosome thus results from the spatial integration of all the peripheral cues, either filtered or amplified, depending on the intrinsic properties of the MT network.

## Acknowledgments

This work has been supported by the European Research Commission (Starting Grant 310472), the Agence Nationale de la Recherche (ANR-12-BSV5-0014-02), the Labex Grenoble Alliance for Integrated Structural Cell Biology and a CTBU Grant from CEA to Gaëlle Letort.

## Materials and Methods

All simulations are performed using Cytosim (www.cytosim.org). The values used of the main parameters are presented in table 1. The motion of elastic fibers surrounded by a viscous fluid (a viscosity of 1 pN.s/μm^2^ was used here (Kole et al., 2005)) is calculated using Langevin dynamics (Nedelec and Foethke, 2007). Microtubules are thus subject to Brownian motion as determined by their size and temperature (k_b_T=4.2 pN.nm).

**Table 1.** Default parameters used in the simulations

|  |  |  |
| --- | --- | --- |
| Microtubules |  |  |
| Rigidity | $k_B T L_p = 25 \text{ pN} \cdot \mu\text{m}^2$ | Persistence length $L_p = 5200 \text{ } \mu\text{m}$ (Gittes et al., 1993) |
| Polymerisation speed | $V_0 = 0.13 \text{ } \mu\text{m/s}$ | (Burakov et al., 2003) |
| Depolymerisation speed | $V_d = 0.27 \text{ } \mu\text{m/s}$ | (Burakov et al., 2003) |
| Rescue rate | $0.064 \text{ s}^{-1}$ | (Burakov et al., 2003) |
| Stall force | $f_s = 5 \text{ pN}$ | (Dogterom and Yurke, 1997) |
| Catastrophe rates | $0.01 \text{ s}^{-1}, 0.04 \text{ s}^{-1}$ | Unloaded catastrophe rate and stalled catastrophe rates. Adapted from (Janson et al., 2003). The unloaded rate is however set higher than observed *in vitro*, to take into account the absence of cytoplasmic obstacles |
| Total tubulin | $M = 10000 \text{ } \mu\text{m}$ | Total tubulin units available in the cell, express as length of MT |
| Centrosome |  |  |
| Radius | $0.5 \text{ } \mu\text{m}$ | Radius of centrosome bead |
| Mobility | $0.03 \text{ } \mu\text{m/pN/min}$ | From (Zhu et al., 2010). In cytosim, the mobility is calculated by setting an effective viscosity of $200 \text{ pN s/}\mu\text{m}^2$ for the bead around which the centrosome is built. This viscosity is only used calculate the mobility of the bead. It does not affect the mobility of MTs. |
| First anchoring stiffness | $k_a = 500 \text{ pN/}\mu\text{m}$ | Stiffness of first link anchoring MT minus ends to centrosome center |
| Second anchoring stiffness | $k_b = 0 \text{ or } 500 \text{ pN/}\mu\text{m}$ | Stiffness of second link anchoring MT to a point on the centrosome periphery |
| Number of MTs | 100 | (Zhu et al., 2010) |
| Dyneins |  |  |
| Motion speed | $V_{\max} = 1.5 \text{ } \mu\text{m/s}$ | Unloaded speed toward the minus end of MTs (Gross et al., 2000) |
| Stall force | $f_{sm} = 1.1 \text{ pN}$ | Stall force of a dynein motor (Gross et al., 2000) |
| Spring stiffness | $k_d = 100 \text{ pN/}\mu\text{m}$ | Stiffness of the link between the dynein motor head and its fixed anchoring position |
| Confinement space |  |  |
| Cell radius | $10 \text{ } \mu\text{m}$ | Radius for the basic circular geometry |
| Confinement stiffness | $k_c = 500 \text{ pN/}\mu\text{m}$ | Confinement strength of MTs inside the space |
| Pinning stiffness | $k_g = 50 \text{ pN/}\mu\text{m}$ | Stiffness of the link that anchors MT plus ends to their contact point at the cell cortex, used to prevent gliding. |

### Microtubules dynamics

MT minus-ends are stably anchored to the centrosome. Plus-ends undergo dynamic instability (Mitchison and Kirschner, 1984) following a two-state model. Each state is implemented in Cytosim as follow:

- polymerization occurs with a speed that is proportional to the free tubulin pool in the cell, and decreases exponentially under opposing force as measured (Dogterom and Yurke, 1997): 
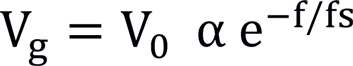

where f is the force component parallel to the axis of the MT, f_s_ the stall force and V_0_ the growth speed parameter, and α is a dimensionless factor in [0,1] representing the fraction of monomer available in the cell. This factor is calculated from the sum of all MT lengths in the cell, divided by the total available tubulin pool, expressed in unit of MT length:

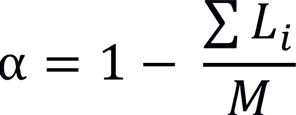
- depolymerisation occurs at a constant speed V_d_.
- catastrophe events occur with a rate τ_c_ that depends on the growth speed of the fiber: 
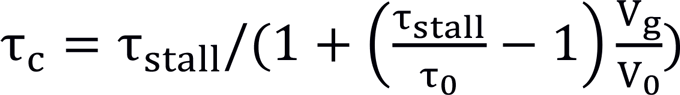

where τ_stall_ is the rate of catastrophe of a stalled microtubules, which is greater than the rate of catastrophe of a free microtubule τ_0_ (Janson et al., 2003).
- rescue events occur at a constant rate τ_r_.

### Microtubules bending elasticity

Microtubules are modeled as semi-flexible polymers (Nedelec and Foethke, 2007), and their buckling thus follows Euler's prediction. If the loading is slow, buckling occurs in the first mode, at a threshold of force, which for a length L and a persistence length L_p_ is:

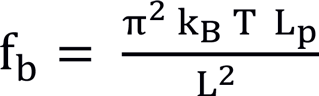

### Centrosome

The centrosome is simulated as an aster with a fixed number of microtubules. All MTs are attached to a small bead of radius R, whose mobility (i.e. the inverse of the drag coefficient) is chosen to match the value of the mobility calculated for the centrosome in (Zhu et al., 2010) to take into account of experimental centrosome motion velocities experimentally measured in (Burakov et al., 2003). The microtubules are anchored at the center of the aster with two Hookean links. The first link of stiffness k_a_ attaches the minus-end of the MTs with the center of the bead. The second link of stiffness k_b_ attaches a distal point on the surface of the bead, with the point of the MT that is located at distance R from the minus end. The number of distal points on the bead is equal to the number of MTs in the aster, and they are distributed regularly around the center of bead, such as to induce an isotropic aster. To allow MTs to pivot, k_b_ is set to zero. In this case, MTs are only constrained by one link, and can freely pivot while their minus-ends remain attached to the center of the bead.

### Confinement

To model the effect of the cortex of the cell on microtubules, the fibers are confined within a fixed geometry. An Hookean force is applied to every microtubule model points that is located outside the confinement geometry:

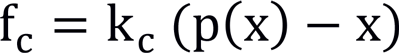

where k_c_ is the spring stiffness and p(x) the projection of the model point x on the edge of the confinement space. This creates a force that is always orthogonal to the edge, thus corresponding to a perfectly slippery edge on which MTs can slide freely. However, in some simulations, the plus end of a MT reaching the edge of the geometry was “pinned” by a spring of stiffness k_g_ acting between the microtubule plus end and the point at which the plus end first reached the edge. When this constraint is added, gliding of microtubules along the cortex is strongly impaired, and the impediment depends on k_g_.

### Motors

A dynein molecule is simulated as a point-like object, that can bind and unbind to microtubules, linked to a fixed position by a spring of stiffness k_d_. This spring represents the anchorage of dyneins either at the cortex or on some vesicle in the cytoplasm. The dynein head moves on a fiber with a speed that depends on the load experienced by the spring:

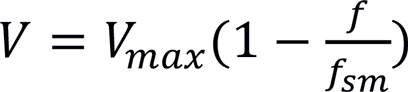

where V_max_ is the speed of a motor without load and f_sm_ is the motor stall force. The value of V_max_ used here is negative representing the fact that dynein head moves toward the minus end of the microtubule.

### Strong cortical motors

Strong cortical motors were added to the simulation to represent the possible effect of local motors associated with proteins such as Par3 in the cortical environment. The particularity of these motors is that they do not unbind unless the microtubule is shrinking. Moreover, these motors stabilize MTs. Specifically, when one or more motors is bound within 0.5 μm of a plus end, the catastrophe rate of this MT plus-end is temporarily set to zero, such that it continues growing.

## Supplementary figures and movies legends

### Figure S1: Effects of pushing force in different cell geometries

Simulations where MT pivoting and gliding are allowed (top) or not allowed (bottom). Snapshots (400 s) for different geometries: ellipse, rectangle, equilateral triangle, acute isosceles triangle and isosceles triangle whose base is the longest side. 15 Trajectories of centrosomes are shown for each geometry. For each trajectory, time is indicated by the color, from blue (0 s) to red (400 s). A black dot indicates the cell gravity center.

### Figure S2: Efficiency of pushing forces when pivoting and gliding are not allowed

(A-C) Trajectories of centrosomes obtained when one parameter is varied: MT rigidity, from 1 to 300 pN.μm^2^; MT unloaded catastrophe rate from 0.01 to 0.06 /s and number of MTs from 15 to 350. (Leftmost two columns) Snapshots (400 s) for 2 different conditions for each varied parameter. (middle column) Multiple trajectories obtained for centrosomes initially positioned along an arc, while the value of the parameter is increasing from left to right. For each trajectory, time is indicated by the color, from blue (0 s) to red (400 s). A black dot indicates the cell gravity center. (right column) Centrosome positioning as a function of one parameter, measured by the distance to the cell center. The dashed lines represent the initial centrosome-center distance (half the confinement radius).

### Movie S1

Simulation for cytoplasmic (left) and cortical (right) dynein distributions in circular geometry. Cell radius is 10 μm. 4000 dynein are uniformly distributed. The system is simulated for 400 s.

### Movie S2

Simulations in different geometries, with cytoplasmic (top) and cortical (bottom) distributions of motors. The geometries are, from left to right: ellipse, rectangle, equilateral triangle, acute isosceles triangle and isosceles triangle whose base is the longest side. The area of all the Geometries is similar, around 314 μm^2^, and contains 4000 dyneins. The system is simulated for 400 s.

### Movie S3

MT network rearrangements in a system dominated by pushing forces. Simulations of MT dynamics with no motors, when MT gliding is allowed (top row) or not (bottom row), and pivoting is allowed (left column) or not (right column). Cell radius is 10 μm. The system is simulated for 400 s.

### Movie S4

Variation of key MT parameters in a system dominated by pushing forces, where MTs are allowed to pivot and glide. The MT rigidity is changed in the top row. The MT catastrophe rate is changed in the middle row, and the number of MTs is changed in the bottom row‥ Cell radius is 10 μm. The system is simulated for 400 s.

### Movie S5

Variation of MT parameters in a system dominated by pushing forces, where MTs are not allowed to pivot or glide. MT rigidity is changed in the top row. MT catastrophe rate is changed in the middle row and MT number is changed in the bottom row. Cell radius is 10 μm. The system is simulated for 400 s.

### Movie S6

Variation of MT parameters in a system where pushing and pulling forces are balanced. The number of cytoplasmic dyneins was set to 100 to match pulling and pushing forces. The MT rigidity is changed in the top left, while pivoting and gliding are allowed. The MT catastrophe rate is changed in the middle left, while pivoting and gliding are allowed. The MT number is changed in the bottom left, while pivoting and gliding are allowed. The MT gliding stiffness is changed in the top right, while pivoting is not allowed. The MT pivoting stiffness is changed in the bottom right, while gliding is not allowed. Cell radius is 10 μm when varying MT rigidity (top left) and 7 μm otherwise. Dynein are shown in green. The system is simulated for 400 s.

### Movie S7

Sensitivity of centering mechanism to external cues.

Simulations with few (300) cytoplasmic dyneins (shown in green) with different numbers of cortical dyneins added position within a 60° crescent at the “bottom” of the cell. In the top row, pivoting and gliding are restricted, and they are allowed in the bottom row. Cell radius is 10 μm. The system is simulated for 400 s.

